# DeepGene: An Efficient Foundation Model for Genomics based on Pan-genome Graph Transformer

**DOI:** 10.1101/2024.04.24.590879

**Authors:** Xiang Zhang, Mingjie Yang, Xunhang Yin, Yining Qian, Fei Sun

## Abstract

Decoding the language of DNA sequences is a fundamental problem in genome research. Mainstream pre-trained models like DNABERT-2 and Nucleotide Transformer have demonstrated remarkable achievements across a spectrum of DNA analysis tasks. Yet, these models still face the pivotal challenge of (1) genetic language diversity, or the capability to capture genetic variations across individuals or populations in the foundation models; (2) model efficiency, specifically how to enhance performance at scalable costs for large-scale genetic foundational models; (3) length extrapolation, or the ability to accurately interpret sequences ranging from short to long within a unified model framework. In response, we introduce DeepGene, a model leveraging Pan-genome and Minigraph representations to encompass the broad diversity of genetic language. DeepGene employs the rotary position embedding to improve the length extrapolation in various genetic analysis tasks. On the 28 tasks in Genome Understanding Evaluation, DeepGene reaches the top position in 9 tasks, second in 5, and achieves the overall best score. DeepGene outperforms other cutting-edge models for its compact model size and superior efficiency in processing sequences of varying lengths. The datasets and source code of DeepGene are available at GitHub (https://github.com/wds-seu/DeepGene).

## 1 Introduction

As a biological language, DNA shares considerable similarities with natural language in many aspects. Presently, the paradigm of modeling human DNA sequences as a natural language holds significant promise [Alharbi and Rashid, 2022, Liu et al., 2020, Zhang et al., 2019]. With the advent of Transformer-based models [Vaswani et al., 2017], numerous large-scale models trained on genetic corpora have emerged. These models employed continuous embeddings to model DNA sequences, demonstrating potential applications across various genomic analysis tasks, including understanding gene regulation [Li et al., 2023], decoding enhancer complexity [Smith et al., 2023], predicting promoters [Lai et al., 2019] and forecasting gene expression [Avsec et al., 2021], etc.

Foundation models for genomics such as DNABERT [Ji et al., 2021], DNABERT-2 [**?**], and Nucleotide Transformer [Dalla-Torre et al., 2023] have achieved state-of-the-art performance across a range of genomics tasks. DNABERT utilized the human reference genome GRCh38.p13 [Guo et al., 2017] for pre-training, adopting K-mer-based sequence tokenization. This approach performed outstandingly in tasks such as promoter prediction and the identification of transcription factor binding sites. As a follow-up work, DNABERT-2 expanded its training dataset to include multi-species genomes and switched to Byte Pair Encoding (BPE) [Sennrich et al., 2016] for DNA segmentation. Additionally, DNABERT-2 established the Genome Understanding Evaluation (GUE) benchmark, demonstrating superior performance over other leading foundation models. The Nucleotide Transformer, pre-trained on a combination of human pan-genome and multi-species genomes, standed out for its use of extensive data volumes and model sizes. Besides, numerous other pre-trained models have emerged [Nguyen et al., 2023, Fishman et al., 2023] in modeling DNA sequences.

Despite significant advancements in foundational models for genetics, several challenges remain: (1) **Reference Genome vs. Pan-genome:** DNABERT and DNABERT-2 were primarily trained on reference genomes. However, the pan-genome concept [Sherman and Salzberg, 2020] captures a broader spectrum of genetic variation within a species than a single reference genome can. It encompasses the diversity of the DNA language, including single nucleotide polymorphisms (SNPs), insertions, deletions, and structural variations across diverse individuals or populations. This breadth of diversity enables a more comprehensive understanding of species-specific genetic variation [Baaijens et al., 2022]. (2) **Performance vs. Costs:** While bigger genomic models tend to exhibit improved performance, the incremental gains begin to diminish as the size of the model and computational resource requirements increase. Finding an optimal balance between performance and cost is critical for the efficient training of genomic models on extensive DNA datasets. (3) **Short Sequence vs. Long Sequence:** The accurate decoding of genomic sequences requires an understanding of extensive contextual information spread across thousands of nucleotides [Zhao et al., 2023]. The maximum input sequence size for DNABERT and Nucleotide Transformer was limited to 512bp and 1000bp, respectively. Gena-LM [Fishman et al., 2023] extended the input sequence length to between 4.5k and 36k base pairs with slide attention [Zaheer et al., 2020], showing superior length extrapolation abilities compared to previous models. However, Gena-LM faced challenges in efficiently analyzing short sequences.

In this study, we introduce DeepGene, a Transformer-based architecture with BPE tokenization for DNA segmentation. The novel contributions lie in (1) DeepGene is trained on a comprehensive dataset comprising 47 ancestrally diverse human pan-genomes. This dataset enables DeepGene to capture the extensive variability of genomic language, providing a more nuanced understanding of genetic diversity; (2) We incorporate the Minigraph representation to efficiently represent the complex graph structures inherent in pan-genomes. This method allows DeepGene to maintain a balance between computational performance and costs, outperforming other cutting-edge genomic models in this respect; (3) Rotary position embedding is applied in DeepGene to accurately learn the relative positions of bases [Huang et al., 2020], which significantly enhances the model’s capability for length extrapolation, improving its performance across both short and long DNA sequences. Experiments on a broad range of datasets and tasks have demonstrated exceptional proficiency in interpreting and predicting genomic sequences, showcasing its potential to contribute valuable insights into the complex language of DNA.

The organization of the paper is as follows: Section 2 offers an in-depth description of the pan-genome dataset utilized as the foundational corpus for pre-training, the implementation of the Minigraph model to depict the graph structures within genomic sequences and the technical details of the DeepGene model. In Section 3, we present the experimental outcomes of DeepGene on GUE among other tasks, engaging in a comparative analysis of its performance relative to other leading models in terms of efficiency and effectiveness. Section 4 provides a summary of our contributions in this research and proposes perspectives for future works.

## 2 Materials and Methods

### 2.1 Pan-genome Dataset

#### 2.1.1 Pan-genome and Minigraph Representation

DNABERT and DNABERT-2 were trained using reference genomes, which were unable to capture the full spectrum of genetic variations. The base model of the Nucleotide Transformer was not only trained on a reference genome but was further trained with samples from thousands of diverse human genomes. While this method significantly expanded the dataset, it also introduced a redundancy rate of 98% among the samples [Dalla-Torre et al., 2023]. This high level of redundancy constrained the ability of the Nucleotide Transformer to learn the diversity of genetic language.

The pan-genome has emerged to encompass population diversity in genomic analyses. It offers a richer source of genetic diversity compared to traditional reference genomes. This enhanced diversity enables a more accurate and efficient understanding of the characteristics of genetic language [Sherman and Salzberg, 2020]. Pan-genome rectifies the biases of reference genomes by storing a representative set of diverse haplotypes and their alignments, leading to the creation of several pan-genomic graph models [Andreace et al., 2023] as de Bruijn graphs [Compeau et al., 2011] and variation graphs [Sirén, 2017]. Due to the natural advantages of graph structures, they are currently the most outstanding in handling large amounts of input data [Eizenga et al., 2020]. The pan-genome graph offers a solution to the data redundancy problem inherent in pan-genomes represented in multiple sequence alignment format. By employing this approach, the model can focus more on the variant parts. De Bruijn graphs utilize k-mers from all haplotypes to construct the graph, where each node represents a k-mer with k-1 overlapping bases between adjacent nodes. In contrast, variation graphs do not enforce a fixed base length for each node; instead, they utilize graphs and path lists to capture input haplotypes. We opt for variation graphs for two key reasons: firstly, De Bruijn graphs exhibit substantial node overlap due to k-mer tokenization, and secondly, the variable node lengths in variation graphs facilitate more refined and flexible tokenization.

The Human Pangenome Reference Consortium recently presented a draft version of the human pan-genome [Liao et al., 2023]. The pan-genome graph in this article is based on the sequencing results of 47 ancestrally diverse humans and was constructed using the methods of Minigraph [Li et al., 2020], Minigraph-Cactus [Armstrong et al., 2020, Hickey et al., 2023], and the Pan-genome Graph Builder (PGGB) [Liao et al., 2023].

As a variation graph, Minigraph asserts the absence of cycles and node overlap within the graph, and its data is presented in the rGFA format, known for its simplicity in processing. Consequently, we choose to utilize this dataset and its Minigraph representation, which is illustrated in Fig. 1.

**Figure 1.**
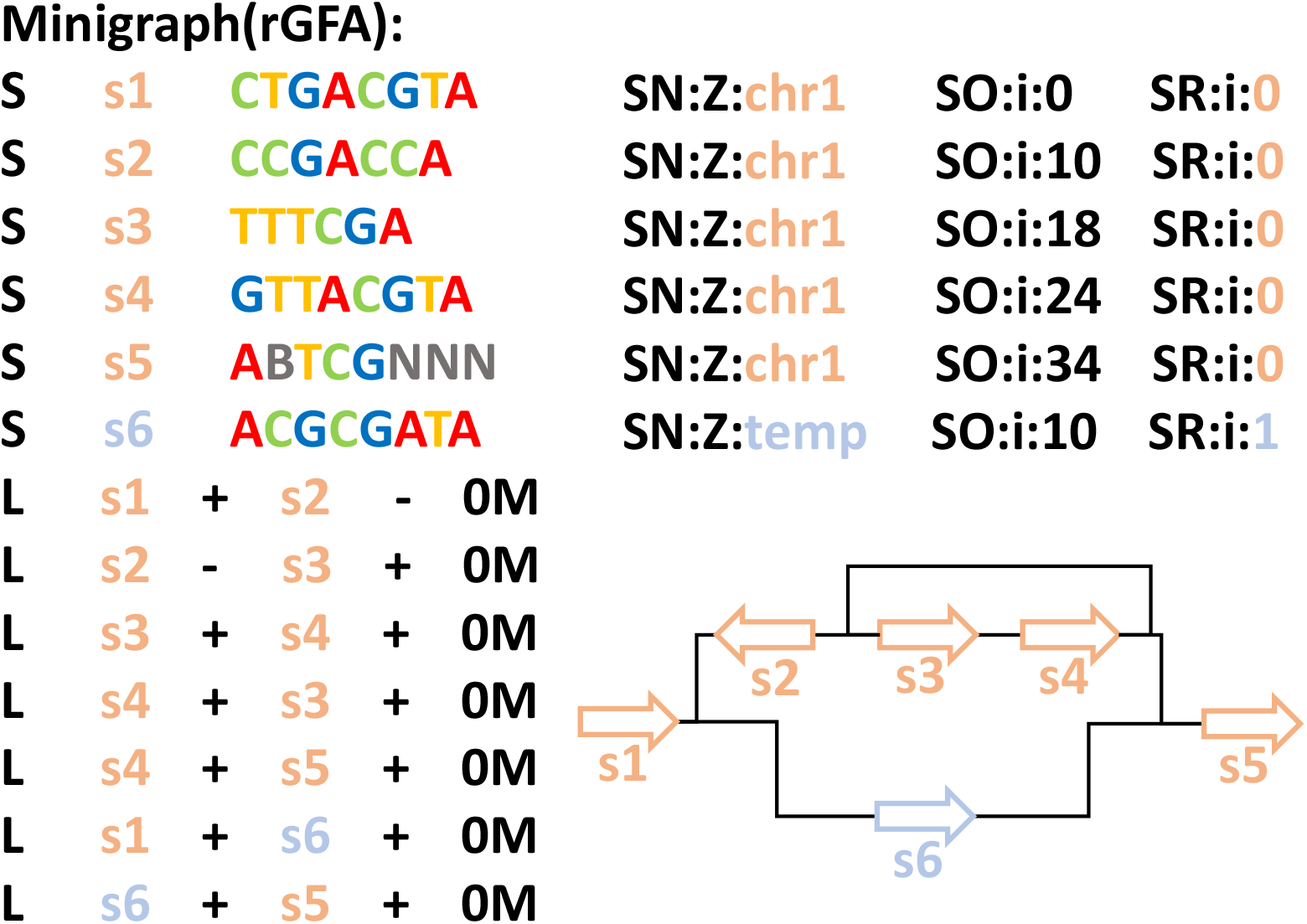
Example of a standard Minigraph representation of pan-genome from genome sequences of two different humans, while the total number of bases is less than 50. The “S” in the figure represents a single node consisting of a segment of nucleotides in a pan-genome. The “L” represents an edge connecting two nodes. “SN” represents the name of sequence from which the node is derived, and “Z” indicates that the content is a printable string, including space. The “SO” means the offset, and “i” indicates that the content is a signed integer. The “SR” is zero if the sequence of a given node is on the reference genome. rGFA disallows overlap between segments. “0M” in the figure means that the length of the overlapping segments between segments is 0. In this example, thick arrows denote segments and thin lines denote links. Different colors indicate different “SR”. The arrow of the segment does not indicate the direction, but indicates whether the segment itself or its reverse complement sequence is used.

The “+” in the line “L” means the segment in the node is the same as the segment in “S”. The “-” in the line “L” means that the segment in the node is the reverse complement segment in “S”. For example, in Fig. 1, the segment in “s1” is “CTGACGTA”. The segment in “s2” is “CCGACCA”. In the first line of “L”, the “s1+”, “CTGACGTA” and the “s2-”, “TGGTCGG”, namely the reverse complement DNA segment connected. There are non-ATCG bases in the nucleotide sequence. Common degenerate base codes are used here, such as “B” for “G” or “T” or “C” and “N” for any base.

The nodes of the minigraph represent the sequences and and are connected by undirected edges. However, the edges on the graph connecting two sequences are ordered because the sequence nodes are differentiated as front and back ends. Therefore, we can consider the direction of the edge as pointing from the back end of one sequence to the front end of another, thus equivalently representing the graph as a directed graph shown as in Fig. 2(a).

**Figure 2.**
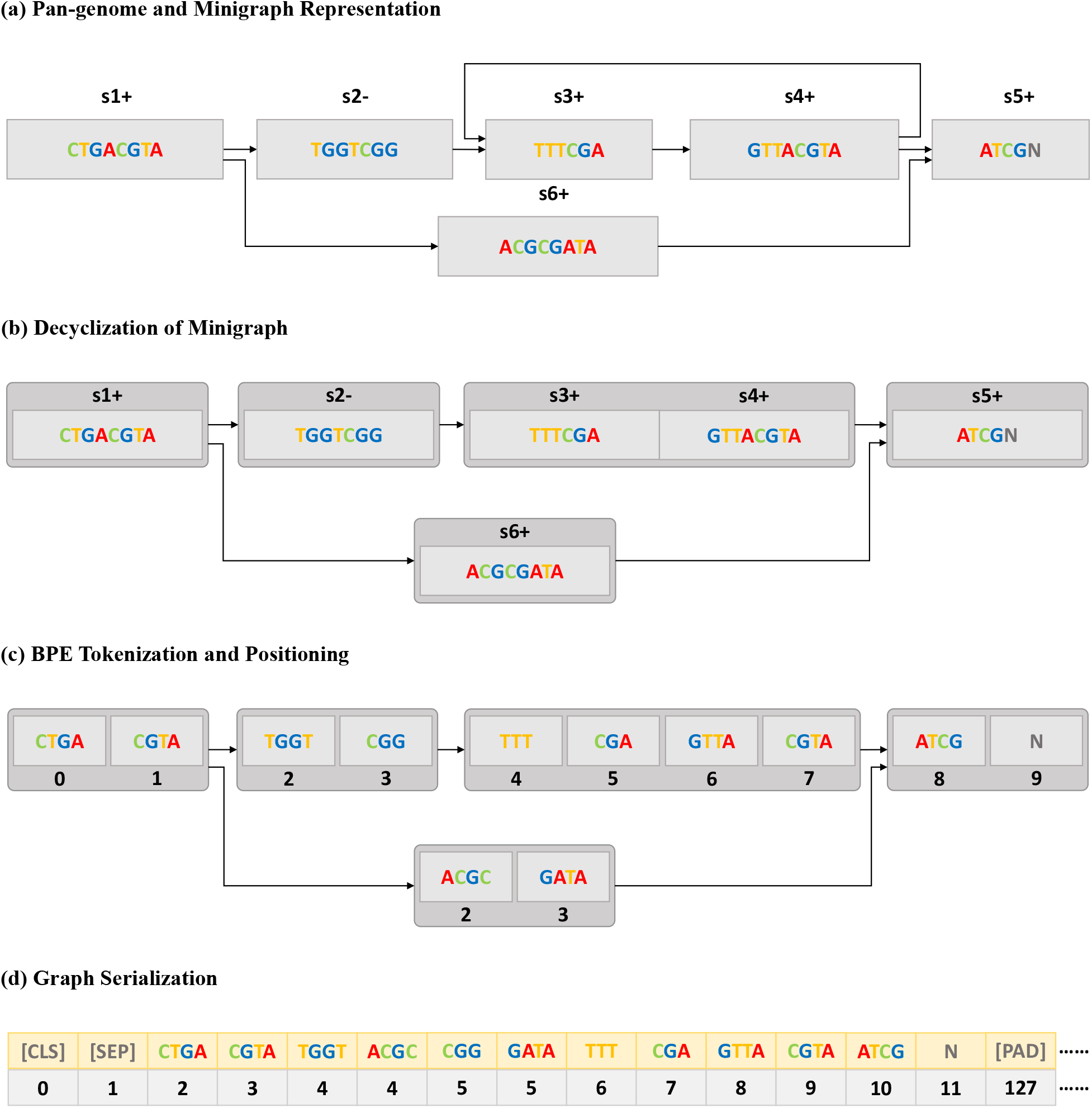
The pipeline of data processing. (a) The result of converting a Minigraph to a directed graph. “s1+” corresponds to the “s1” segment, “s2-” is the reverse complement of the “s2” segment; (b) Decyclization with the strong connectivity algorithm; (c) BPE tokenization and adding positional information. (d) Serialize the graph so that it can be put into Transformer-based model training.

#### 2.1.2 Decyclization of Minigraph

Although many sources claim that the graph in the rGFA file of Minigraph is a directed acyclic graph (DAG) [Andreace et al., 2023], we still found a small number of cycles in the graph during the experiment. The position information of each node will be uncertain without removing them, which will be illustrated in detail below. In response to this situation, we used the strongly connected components algorithm to eliminate the cycles and obtained a directed acyclic graph (DAG) shown as in Fig. 2(b).

#### 2.1.3 BPE Tokenization and Positioning

Due to the possibility that a node contains thousands of base pairs, it is not feasible to consider a single node as the smallest processing unit in the pre-training of the DeepGene model. The DNABERT-2 made the ablation that the BPE is better than most tokenization methods on DNA sequence. Therefore, we utilize BPE for tokenization, splitting each node in minigraph into multiple tokens. Each token will have a specified positional information in the node, which is represented as follows.

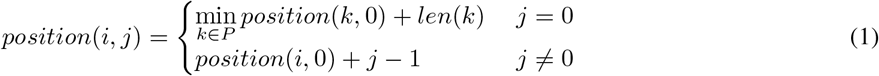

The *position*(*i, j*) refers to the positional information of the *j*-th token in node *i*, and *len*(*k*) denotes the total number of tokens in node *k. P* is the set of predecessor nodes of node *i*. It is important to note that if a node does not have a predecessor node, then its *position*(*i*, 0) is 0. This is illustrated in Fig. 2(c).

#### 2.1.4 Graph Serialization

It is known that Transformer-based models impose a maximum input length constraint, making it infeasible to input an entire Minigraph into the model as a single sample. Hence, we sort all tokens in the connected Minigraph by their position and partition the Minigraph into subgraphs, ensuring that each subgraph contains no more than 126 tokens. Special tokens such as [CLS], [SEP], and [PAD] are then appended to each subgraph until it reaches a length of 128 tokens. Positional information is assigned to these tokens, with the [CLS] position designated as 0, the [SEP] position as 1, and subsequent [PAD] positions as 127. Additionally, the positions of all other tokens are adjusted collectively, starting from 2. The incorporation of padding and positional encoding for special tokens ensures all sequences within a batch are complete and have the same length. This is illustrated in Fig. 2(d).

#### 2.1.5 Dataset Statistics

The pan-genome utilized as the pre-training corpus for DeepGene comprises more than 5 billion total base pairs, divided into over 1 billion tokens. Ultimately, approximately 10 million subgraphs are generated by padding special tokens alongside sequence tokens to ensure each subgraph consists of a uniform 128 tokens. Each subgraph is then individually pre-trained as a separate sample.

### 2.2 DeepGene Model

Existing foundation models for genomes often encounter difficulties in handling long sequences due to their limited ability to directly capture relationships between distant genomic tokens. To improve the length extrapolation capability of the model, we integrate Rotary Position Embedding (RoPE in short) [Su et al., 2024] into DeepGene. RoPE addresses this issue by introducing rotation operations. These operations rotate the vector space of positional encodings, making relationships between distant genomic tokens clearer in the vector space.

#### 2.2.1 Rotary Position Embedding

RoPE is an enhancement to the traditional positional encoding used in Transformer-based models. In contrast to conventional positional encoding, which merely adds fixed sinusoidal or learned positional embeddings to word embeddings, RoPE introduces a rotation mechanism to more effectively capture positional relationships within the input sequence. It initially processes the *Q* and *K* vectors and incorporates absolute positional information into *Q* and *K*. Specifically, after mapping the input to *Q* and *K*, the *Q* and *K* vectors are rotated by multiplying with a rotary matrix 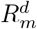. The rotary matrix 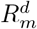 contains positional sequence information. The representation of 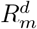 is as follows:

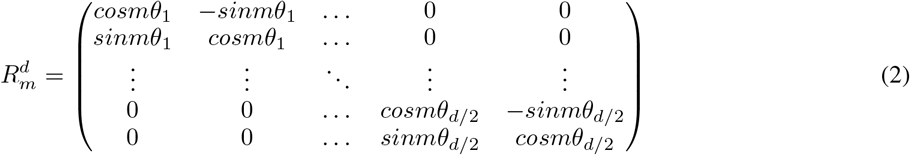

The symbol *m* denotes the position of a token, *d* represents the dimension of the hidden state, and *θ* is a constant. Following the rotary operation, incorporating the absolute positional information of *Q* and *K* into the attention calculation will fuse the attention matrix with relative positional information through the dot product of *Q* and *K*. This operation effectively achieves relative positional encoding in an absolute positional encoding manner. As shown in the following formula, the dot product obtained by multiplying the input vectors at positions *m* and *n* by the rotary matrix automatically incorporates the relative positional information of *m* and *n* into self-attention.

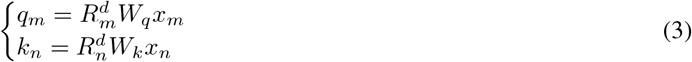

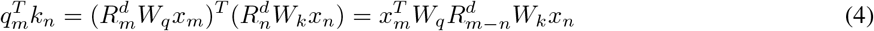

The core idea behind rotary positional encoding is to represent relative positional information through absolute positions in the attention mechanism, enabling the model to discern the relative positions between gene sequence segments rather than solely absolute positions. Experimental results indicate that relative positional encoding yields superior results compared to absolute positional encoding. Moreover, it enhances the model with length extrapolation capability, meaning that despite being pre-trained on short sequences (e.g., 128bp), the model exhibits comparable performance on both short and long sequences.

#### 2.2.2 Model Pre-training

The model architecture is shown in Fig. 3. During the pre-training process, the DeepGene model first processes graph data into sequences and then applies masking. After embedding, the input undergoes a RoPE-based multi-head attention module. The detail of this module is shown in the figure and illustrated in the rotary position embedding mentioned above. Subsequently, the model applies softmax to the attention matrix *A*, multiplies it with *V*, and proceeds with feedforward and layer norm operations. Ultimately, this process yields the representation of the gene sequence. Similar to BERT, the pre-training task adopts the classic MLM task to predict tokens that have been chosen randomly. The proportion of tokens chosen is 15%. Among these tokens, 80% of them are masked, 10% are randomly replaced and 10% remain unchanged.

**Figure 3.**
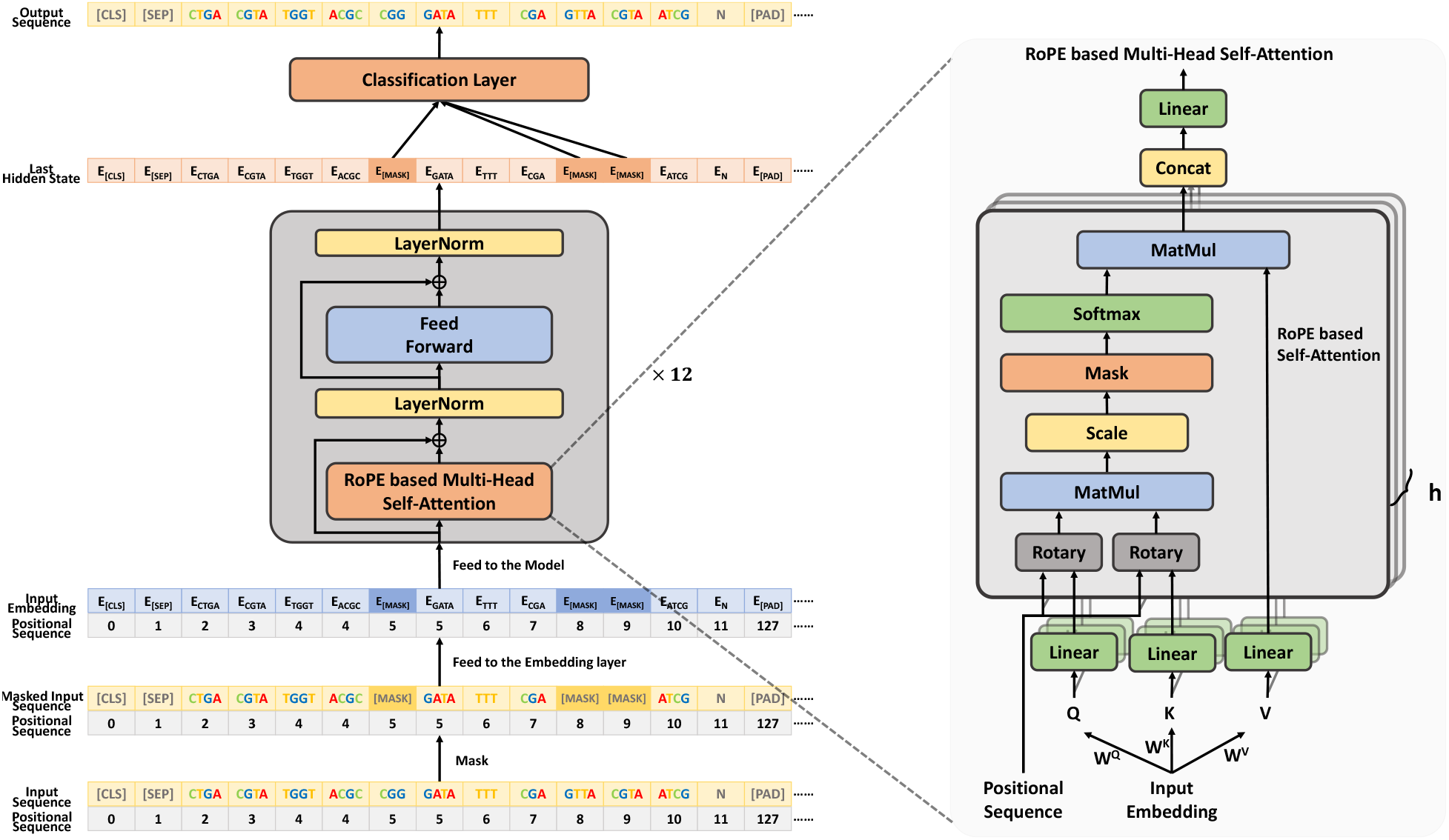
Details of the architecture and characteristics of the DeepGene model. The sequence is tokenized by BPE algorithm as the input. The input passes the embedding layer and is fed into 12 Rope Transformer blocks to obtain the relative position information. The MLM method, which predicts the masked token is used to pre-train the model.

**Figure 4.**
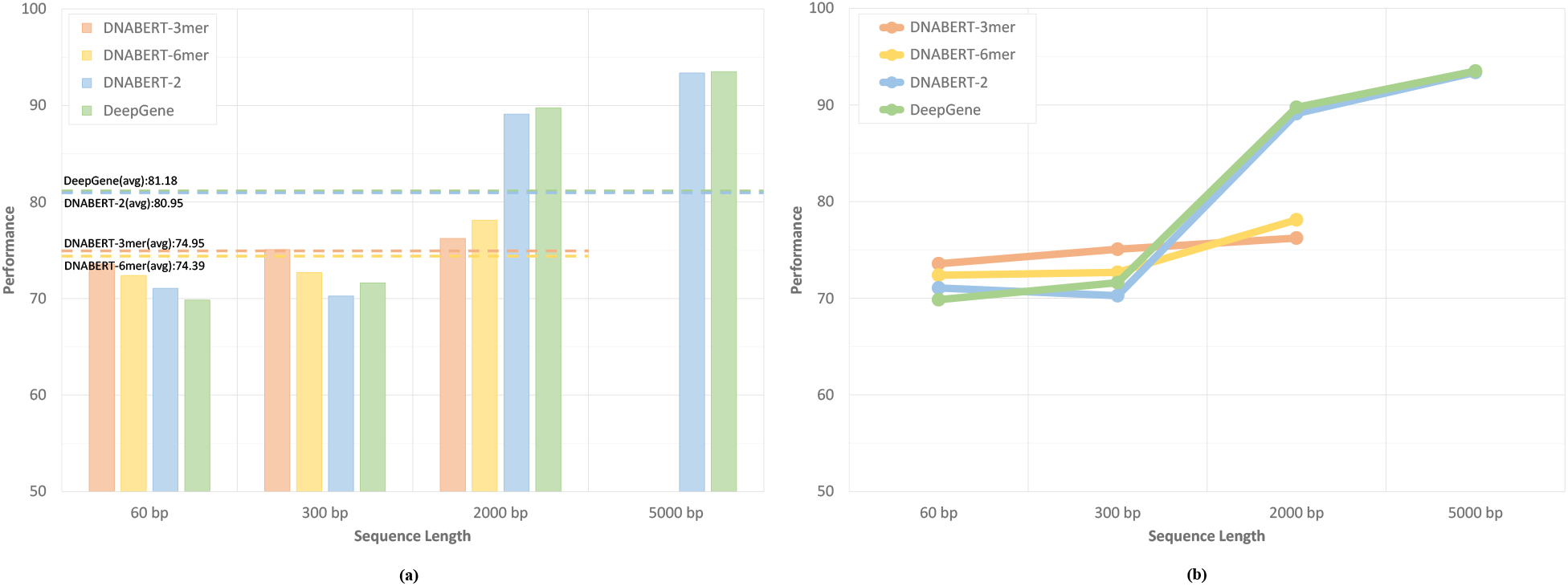
(a) The result of DeepGene, DNABERT and DNABERT-2 on the promoter detection data in different lengths. It is obvious that with the length increasing, the performance of the DeepGene and DNABERT-2 soar while DNABERT increases slowly. On the 2000bp task, DeepGene achieves more than 10% scores compared with DNABERT 3-mer and 6-mer, and it is also slightly better than DNABERT-2. On the 5000bp task, the f1 score obtained by our model is also the highest. (b) The performance trend of each foundation model.

The pre-training task is conducted for 20 epochs, with a batch size of 2,048. During the first 12 epochs, we implement a warm-up phase, where the learning rate linearly increased from 0 to 4 *×* 10^−4^. For the subsequent 8 epochs, the learning rate linearly decreased from 4 *×* 10^−4^ to 1.33 *×* 10^−4^. We utilize 4 A100 (40G) GPUs for pre-training and completed the process in only four days, significantly faster than other models. We employ a model architecture similar to the BERT base, consisting of 12 layers with a hidden layer dimension of 768.

#### 2.2.3 Model Fine-tuning

After pre-training on graph-modeled pan-genome data, we fine-tune DeepGene across various downstream tasks using sequence-based genomic data, since the downstream tasks and baselines are all sequence-based. Given the relatively modest model size, we opt for full-parameter fine-tuning directly. The fine-tuning of DeepGene encompasses not only short gene sequences in downstream tasks but also longer sequences to validate the model’s substantial length extrapolation capabilities. We utilize the hidden layer vector of the [CLS] token at a sequence’s outset to represent the sequence. Employing a Multi-Layer Perceptron (MLP) as the classification head, we conduct pertinent classification tasks.

## 3 Experiments and Results

### 3.1 Baselines

**DNABERT** was the pioneering pre-trained model designed for genome sequences. This model was pre-trained on human genomes, consisting of four variants named DNABERT (3-mer), DNABERT (4-mer), DNABERT (5-mer), and DNABERT (6-mer). They were different in vocabulary size depending on the choice of k-mer.

**Nucleotide Transformer** had a much larger size of model parameters and datasets. It used pan-genome dataset. It had achieved state-of-art performance in many DNA analysis tasks. It had four variants too: NT-500M-human, NT-500M-1000g, NT-2500M-1000g and NT-2500M-multi. The first model was pre-trained on human reference genome with a model size of 500M. The second was pre-trained on the pan-genome of 3,202 humans with a model size of 500M. The third was pre-trained on the pan-genome of 3,202 humans with a model size of 2500M. The last one was pre-trained on 850 genomes of multi-species with a model size of 2500M.

**DNABERT-2** was an improved model based on DNABERT. It was pre-trained on a multi-species genome dataset. The model used BPE as the tokenization method, utilizing many computational methods like relative position embedding. It achieved great performance on many DNA analysis tasks.

### 3.2 Experiement Datasets and Settings

We assess DeepGene across two distinct task categories. The first involves Genomic Understanding and Evaluation ^1^ (GUE in short), introduced by DNABERT-2, emphasizing the capability to effectively address comprehensive downstream tasks in gene analysis. The second category is Long Promoter Detection (LPD in short), which specifically evaluates the model’s proficiency in comprehending long DNA sequences.

#### 3.2.1 Genomic Understanding and Evaluation (GUE)

To verify that DeepGene, as a foundation model, has learned representations of DNA sequences from the pre-training corpus, we conduct a series of downstream task experiments to evaluate its performance across different downstream tasks in GUE as proposed by DNABERT-2. The GUE assessment criteria encompassed a range of diverse classification tasks. It included seven major categories and a total of 28 subtasks. These tasks covered not only the human genome but also the genomes of various species such as yeast, mice, and viruses. Furthermore, the tasks included both binary classification and multi-class classification. The sequence lengths ranged from 70bp to 1000bp. This comprehensive setup allowed for a thorough evaluation of the model’s ability to represent different genes. Except for virus-related tasks, which utilized the f1-score metric, all other tasks employed Matthews’s Correlation Coefficient (MCC) for evaluation. During fine-tuning, we employ full-parameter fine-tuning with a uniform batch size of 32. The weight decay is 0.01. We initiate a warm-up for the first 50 steps of fine-tuning, linearly increasing the learning rate from 0 to 3 *×* 10^−5^, and then maintains a constant learning rate for the remainder of the fine-tuning process. Every 200 steps during fine-tuning, the model is evaluated, and the best-performing model on the validation set across all evaluations is selected for testing. Due to variations in the data scale for different downstream tasks, the total number of fine-tuning steps differs for each task, as indicated in Tabel. 1.

**Table 1:** The number of training steps for the following tasks. Epigenetic Marks Prediction (EMP), Transcription Factor Prediction on the Human genome and the Mouse genome (TF-H and TF-M), Covid Variants Classification (CVC), TATA dataset of Promoter Detection (PD-TATA), non-TATA and all datasets of Promoter Detection (PD-other), TATA dataset of Core Promoter Detection (CPD-TATA), non-TATA and all datasets of Core Promoter Detection (CPD-other), and Splice Site Prediction (SSP).

|  | EMP | TF-M | CVC | TF-H | PD-TATA | PD-other | CPD-TATA | CPD-other | SSP |
| --- | --- | --- | --- | --- | --- | --- | --- | --- | --- |
| Num. Epochs | 5 | < 2k steps | 10 | 5 | 10 | 5 | 10 | 5 | 5 |

#### 3.2.2 Long Promoter Detection (LPD)

To assess DeepGene’s proficiency in understanding lengthy sequences, we opt for the classic downstream task of promoter recognition to conduct a length extrapolation experiment. We employ the methodology utilized by Gena-LM to acquire data relevant to promoter recognition downstream tasks. The primary data source is EPDNew [Dreos et al., 2013]. We utilize the EPDNew select tool to retrieve human promoter sequences (hg38). To effectively illustrate the impact of varying sequence lengths on the model’s performance, we select three distinct lengths of promoter sequence fragments varying from 60bp (short sequence), 300bp (medium sequence), 2000bp (long sequence) to 5000bp (extra long sequence) for experimentation. Subsequently, we utilize the Gena-LM approach to generate corresponding negative examples and divide the data into training, validation, and testing sets in a 3:1:1 ratio.

The specific settings of the obtained promoter sequence dataset are detailed in Tabel. 2.

**Table 2:** The illustration of promoter detection dataset. The data in different lengths are sampled from different positions on the stable sequences.

| Length | Begin | End | Train / Dev/ Test |
| --- | --- | --- | --- |
| 60bp | -29bp | 30bp |  |
| 300bp | -249bp | 50bp | 35518/11839/11839 |
| 2000bp | -1000bp | 999bp |  |
| 5000bp | -2000bp | 2999bp |  |

The maximum length of the sequences during DNABERT pre-training is 512 tokens. Since it uses the overlapping k-mer tokenization, the maximum number of nucleotides is 512-k+1, which is approximately 510bp. The maximum length of the sequences during pre-training of DNABERT-2 and our model is 128 tokens. Since the BPE tokenization without overlapping is used, the average number of bases represented by a token is about 5.5, so the maximum number of nucleotides modeled is about 700, which is roughly the same level as DNABERT. The maximum number of nucleotides modeled during Nucleotide Transformer pre-training is 6000bp, which is much larger than other models. Apparently it has the ability to model long sequences. However, due to our computing power limitations, there is no way to perform full parameter fine-tuning on it. So we can only give up comparing with it. Fortunately, DNABERT-2 has used Nucleotide Transformer to do long sequence tasks, and the final results show that DNABERT-2 has better length modeling capabilities than Nucleotide Transformer. To sum up, we finally chose to compare with DNABERT-2 and DNABERT. Comparison under similar pre-training sequence lengths and model sizes can better reflect the length extrapolation capability of different positional encodings. DNABERT offered multiple versions based on different tokenizations, and the choice of k-mer may significantly impact its length extrapolation capability. In our comparison experiment, we included two versions of DNABERT: 3-mer and 6-mer, representing the smallest and largest k-mer versions, respectively.

We use the same classification head during the evaluation to ensure a fair assessment of the models’ ability to model sequences of different lengths. Specifically, we utilized the feature vector of the [CLS] token in the last hidden layer of the pre-trained model as the representation for the entire sequence. Subsequently, we connected an MLP classification head for binary classification tasks, with the f1-score serving as the evaluation metric. During fine-tuning, we adopted full-parameter fine-tuning with a consistent batch size of 128 and a weight decay of 0.01. In the initial 50 steps of fine-tuning, we implemented warm-up, linearly increasing the learning rate from 0 to 3 × 10^−5^, and then maintained a constant learning rate for the remainder of the fine-tuning process. For sequences of varying lengths (short, medium, and long), we conducted fine-tuning for three epochs, evaluating the model every half epoch. The best-performing model on the validation set was selected for final testing.

### 3.3 The Performance and Costs Evaluation on GUE

We categorized the downstream tasks by type, as shown in Tabel. 3. The table shows the results for all 28 subtasks in 7 major categories.

**Table 3:**
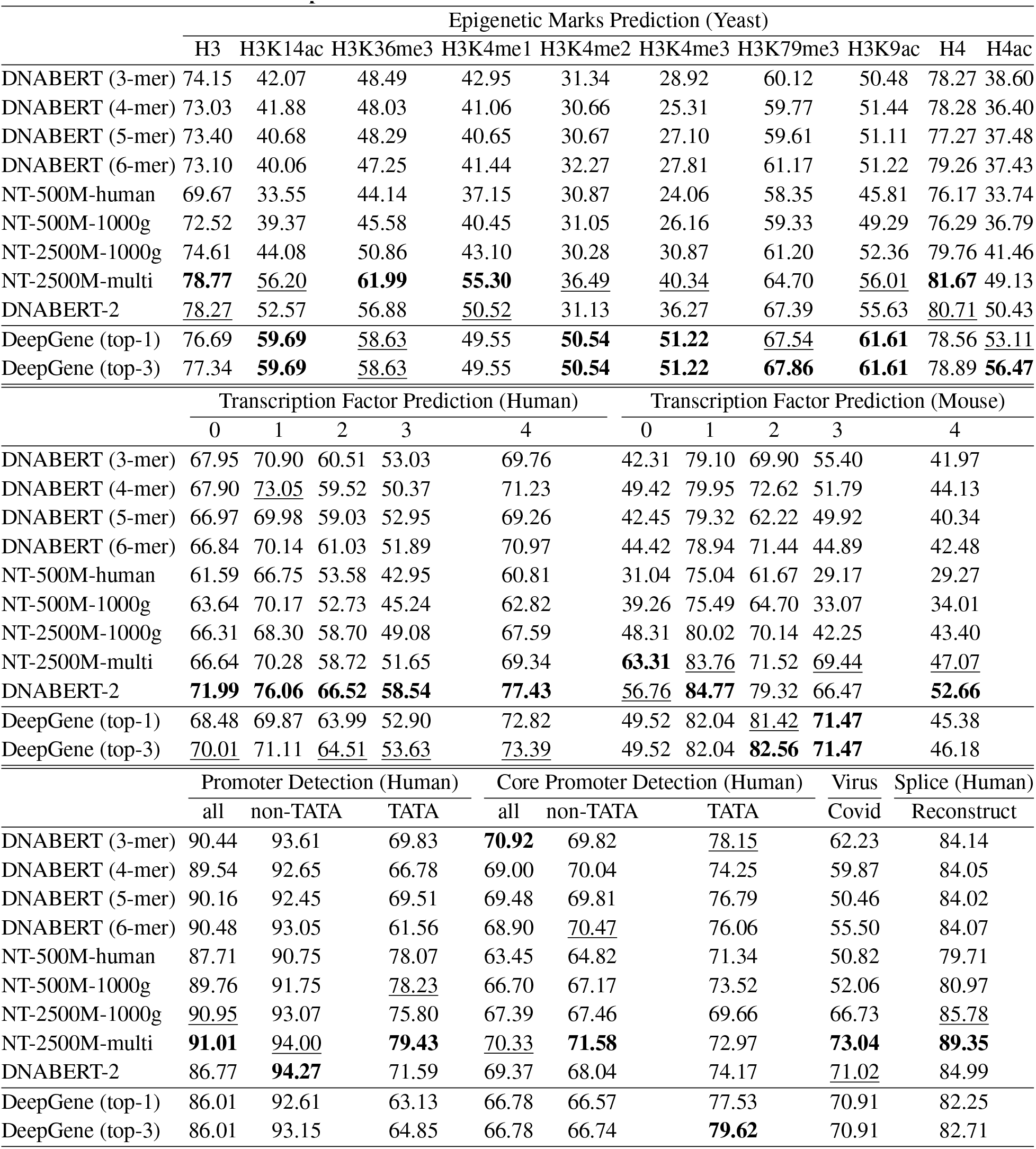
The performance of different models on the GUE benchmark.

|  | Epigenetic Marks Prediction (Yeast) |  |  |  |  |  |  |  |  |  |
| --- | --- | --- | --- | --- | --- | --- | --- | --- | --- | --- |
|  | H3 | H3K14ac | H3K36me3 | H3K4me1 | H3K4me2 | H3K4me3 | H3K79me3 | H3K9ac | H4 | H4ac |
| DNABERT (3-mer) | 74.15 | 42.07 | 48.49 | 42.95 | 31.34 | 28.92 | 60.12 | 50.48 | 78.27 | 38.60 |
| DNABERT (4-mer) | 73.03 | 41.88 | 48.03 | 41.06 | 30.66 | 25.31 | 59.77 | 51.44 | 78.28 | 36.40 |
| DNABERT (5-mer) | 73.40 | 40.68 | 48.29 | 40.65 | 30.67 | 27.10 | 59.61 | 51.11 | 77.27 | 37.48 |
| DNABERT (6-mer) | 73.10 | 40.06 | 47.25 | 41.44 | 32.27 | 27.81 | 61.17 | 51.22 | 79.26 | 37.43 |
| NT-500M-human | 69.67 | 33.55 | 44.14 | 37.15 | 30.87 | 24.06 | 58.35 | 45.81 | 76.17 | 33.74 |
| NT-500M-1000g | 72.52 | 39.37 | 45.58 | 40.45 | 31.05 | 26.16 | 59.33 | 49.29 | 76.29 | 36.79 |
| NT-2500M-1000g | 74.61 | 44.08 | 50.86 | 43.10 | 30.28 | 30.87 | 61.20 | 52.36 | 79.76 | 41.46 |
| NT-2500M-multi | **78.77** | <u>56.20</u> | **61.99** | **55.30** | <u>36.49</u> | <u>40.34</u> | 64.70 | <u>56.01</u> | **81.67** | 49.13 |
| DNABERT-2 | <u>78.27</u> | 52.57 | 56.88 | <u>50.52</u> | 31.13 | 36.27 | 67.39 | 55.63 | <u>80.71</u> | 50.43 |
| DeepGene (top-1) | 76.69 | **59.69** | <u>58.63</u> | 49.55 | **50.54** | **51.22** | <u>67.54</u> | **61.61** | 78.56 | <u>53.11</u> |
| DeepGene (top-3) | 77.34 | **59.69** | <u>58.63</u> | 49.55 | **50.54** | **51.22** | **67.86** | **61.61** | 78.89 | **56.47** |
|  | Transcription Factor Prediction (Human) |  |  |  |  | Transcription Factor Prediction (Mouse) |  |  |  |  |
|  | 0 | 1 | 2 | 3 | 4 | 0 | 1 | 2 | 3 | 4 |
| DNABERT (3-mer) | 67.95 | 70.90 | 60.51 | 53.03 | 69.76 | 42.31 | 79.10 | 69.90 | 55.40 | 41.97 |
| DNABERT (4-mer) | 67.90 | <u>73.05</u> | 59.52 | 50.37 | 71.23 | 49.42 | 79.95 | 72.62 | 51.79 | 44.13 |
| DNABERT (5-mer) | 66.97 | 69.98 | 59.03 | 52.95 | 69.26 | 42.45 | 79.32 | 62.22 | 49.92 | 40.34 |
| DNABERT (6-mer) | 66.84 | 70.14 | 61.03 | 51.89 | 70.97 | 44.42 | 78.94 | 71.44 | 44.89 | 42.48 |
| NT-500M-human | 61.59 | 66.75 | 53.58 | 42.95 | 60.81 | 31.04 | 75.04 | 61.67 | 29.17 | 29.27 |
| NT-500M-1000g | 63.64 | 70.17 | 52.73 | 45.24 | 62.82 | 39.26 | 75.49 | 64.70 | 33.07 | 34.01 |
| NT-2500M-1000g | 66.31 | 68.30 | 58.70 | 49.08 | 67.59 | 48.31 | 80.02 | 70.14 | 42.25 | 43.40 |
| NT-2500M-multi | 66.64 | 70.28 | 58.72 | 51.65 | 69.34 | **63.31** | <u>83.76</u> | 71.52 | <u>69.44</u> | <u>47.07</u> |
| DNABERT-2 | **71.99** | **76.06** | **66.52** | **58.54** | **77.43** | <u>56.76</u> | **84.77** | 79.32 | 66.47 | **52.66** |
| DeepGene (top-1) | 68.48 | 69.87 | 63.99 | 52.90 | 72.82 | 49.52 | 82.04 | <u>81.42</u> | **71.47** | 45.38 |
| DeepGene (top-3) | <u>70.01</u> | 71.11 | <u>64.51</u> | <u>53.63</u> | <u>73.39</u> | 49.52 | 82.04 | **82.56** | **71.47** | 46.18 |
|  | Promoter Detection (Human) |  |  | Core Promoter Detection (Human) |  |  | Virus | Splice (Human) |  |  |
|  | all | non-TATA | TATA | all | non-TATA | TATA | Covid | Reconstruct |  |  |
| DNABERT (3-mer) | 90.44 | 93.61 | 69.83 | **70.92** | 69.82 | <u>78.15</u> | 62.23 | 84.14 |  |  |
| DNABERT (4-mer) | 89.54 | 92.65 | 66.78 | 69.00 | 70.04 | 74.25 | 59.87 | 84.05 |  |  |
| DNABERT (5-mer) | 90.16 | 92.45 | 69.51 | 69.48 | 69.81 | 76.79 | 50.46 | 84.02 |  |  |
| DNABERT (6-mer) | 90.48 | 93.05 | 61.56 | 68.90 | <u>70.47</u> | 76.06 | 55.50 | 84.07 |  |  |
| NT-500M-human | 87.71 | 90.75 | 78.07 | 63.45 | 64.82 | 71.34 | 50.82 | 79.71 |  |  |
| NT-500M-1000g | 89.76 | 91.75 | <u>78.23</u> | 66.70 | 67.17 | 73.52 | 52.06 | 80.97 |  |  |
| NT-2500M-1000g | <u>90.95</u> | 93.07 | 75.80 | 67.39 | 67.46 | 69.66 | 66.73 | <u>85.78</u> |  |  |
| NT-2500M-multi | **91.01** | <u>94.00</u> | **79.43** | <u>70.33</u> | **71.58** | 72.97 | **73.04** | **89.35** |  |  |
| DNABERT-2 | 86.77 | **94.27** | 71.59 | 69.37 | 68.04 | 74.17 | <u>71.02</u> | 84.99 |  |  |
| DeepGene (top-1) | 86.01 | 92.61 | 63.13 | 66.78 | 66.57 | 77.53 | 70.91 | 82.25 |  |  |
| DeepGene (top-3) | 86.01 | 93.15 | 64.85 | 66.78 | 66.74 | **79.62** | 70.91 | 82.71 |  |  |

In the table, we show the performance of DeepGene with two versions: DeepGene (top-1) and DeepGene (top-3). For DeepGene (top-1), the model performance stands for the testing performance of the fine-tuned DeepGene model that performs best on the validation set. While for DeepGene (top-3), we used another strategy for model selection in testing. Top-3 fine-tuned DeepGene (each with 200 steps of fine-tuning) models on the validation set were all selected, and the results represented the best testing performance among these top-3 models.

Among ten subtasks on Yeast sequences, our model achieved first place in 6 subtasks including H3K14a, H3K4me2, H3K4me3, H3K79me, H3K9ac, H4ac, and second place in H3K36me3. In addition, in subtasks such as H3K4me2 and H3K4me, DeepGene outperformed the best baseline model on MCC by more than 10%. Among the five subtasks on Mice, our model also achieves two first places, which also surpasses the current best baseline model by 3% and 5% in MCC. In subtasks on Human and virus, our model win one first place and four second places.

Despite being trained solely on human pan-genome graphs, DeepGene demonstrates a remarkable ability to generalize in modeling gene sequences across multiple species. We suppose that the linguistic diversity inherent in human pan-genome graphs facilitates DeepGene’s transfer of language knowledge, aiding in the comprehension of DNA sequences from other species.

However, we have also observed that while DeepGene demonstrates outstanding performance on yeast and mouse sequences, its performance on tasks related to human sequences is relatively moderate. We attribute this to the following factors: GUE subtasks involving human sequences are based on the human reference genome, whereas our model is trained on pan-genome graphs. This leads to a data bias between the pre-training and testing sequences on the human genome. This bias acts as a double-edged sword: it lowers DeepGene’s performance on tasks related to humans but enhances its performance on tasks involving other species, where the understanding of greater language diversity is required.

By averaging the results of all 28 downstream subtasks and considering various factors such as the scale of the model and data, we present a comprehensive evaluation table in Tabel. 4 to illustrate the balance between performance and cost for each model. DeepGene stands as the smallest among all mainstream foundation models. During inference, it requires the least amount of computation. During pre-training, it has the smallest total number of training tokens, and its pre-training corpus is slightly larger than DNABERT’s but considerably smaller than that of DNABERT-2 and Nucleotide Transformer. Despite this, DeepGene (top-3) achieves the highest average score among all foundation models, surpassing the suboptimal NT model by 0.5% while reducing the model size by 95%. Overall, DeepGene achieves 9 first-place and 5 second-place rankings across all 28 subtasks. This strongly suggests that DeepGene effectively leverages the presently limited corpus of human pan-genome graphs to strike a superior balance between performance and costs.

**Table 4:** The scale of model and data of each mainstream model and their average performance. The comparative table shows that our model achieves great performance on par with current mainstream models while using smaller parameters and pre-trained data. Additionally, the computational cost of our model during inference is smaller. To assess the computational cost, we consider Floating Point Operations (FLOPs), which represents the total number of multiplication and addition operations during a forward pass. We measure FLOPs using genome sequences with a length of 500 during inference. The FlOPs in the table is relative FLOPs compared to DeepGene.

| Model | Num. Params. ↓ | FLOPs ↓ | Trn. Tokens | Pre-Trn. Datasize (bp) | Num. Top-2 ↑ | Ave. Scores ↑ |
| --- | --- | --- | --- | --- | --- | --- |
| DNABERT (3-mer) | 86M | 4.35 | 122B | 3B | 1/1 | 61.62 |
| DNABERT (4-mer) | 86M | 4.34 | 122B | 3B | 0/1 | 61.14 |
| DNABERT (5-mer) | 87M | 4.34 | 122B | 3B | 0/0 | 60.05 |
| DNABERT (6-mer) | 89M | 4.32 | 122B | 3B | 0/1 | 60.51 |
| NT-500M-human | 480M | 4.24 | 50B | 3B | 0/0 | 55.43 |
| NT-500M-1000g | 480M | 4.24 | 50B | 115B | 0/1 | 58.23 |
| NT-2500M-1000g | 2537M | 25.86 | 300B | 115B | 0/2 | 61.41 |
| NT-2500M-multi | 2537M | 25.86 | 300B | 174B | 10/9 | <u>66.93</u> |
| DNABERT-2 | 117M | 1.33 | 262B | 32.5B | 8/5 | 66.80 |
| DeepGene (top-1) | 85M | 1.00 | 26B | 5B | 5/4 | 66.82 |
| DeepGene (top-3) | 85M | 1.00 | 26B | 5B | 9/5 | **67.43** |

### 3.4 The Length Extrapolation Evaluation on LPD

We conducted LPD evaluations on DNABERT, DNABERT2, and DeepGene, with the results shown in Figure 5(a). Across four different sequence lengths, DeepGene averagely outperforms other foundation models, particularly on longer sequences. While its performance on short and medium sequences is moderate, DeepGene significantly surpasses DNABERT by more than 11% on long sequences of 2000bp. In the case of the extra-long task of 5000bp sequences, DNABERT-3mer and -6mer are unable to fulfill the task due to excessive computational resource requirements. DNABERT-2 is slightly outperformed by DeepGene on both long and extra-long sequence tasks.

The trend chart in Figure 5(b) offers further insights into how performance evolves with sequence length. As the length gradually increases, the performance of each model demonstrates continuous improvement, highlighting the significant contribution of longer sequences to promoter recognition tasks. However, the improvement in DNABERT’s performance plateaus at 2000bp, whereas DNABERT-2 and our model show substantial enhancements. Notably, even at 5000bp, a notable improvement is still observed. This underscores the effectiveness of relative positional encoding in modeling relatively longer sequences compared to absolute positional encoding. Particularly, when the sequence length exceeds the maximum encountered during pre-training, the disparity becomes more pronounced, showcasing the model’s remarkable length extrapolation capability. Furthermore, considering that our rotary position embedding does not require additional parameters like ALiBi in DNABERT-2, our approach holds additional advantages.

### 3.5 Discussion

While DeepGene has demonstrated its efficacy as a foundation model for genomics, it still exhibits limitations on certain tasks. Our experiments offer insights for further harnessing the potential of pan-genome-graph-based models like DeepGene. Firstly, while the BPE tokenization method improves model performance, it relies on statistics from reference genomes, highlighting the need for a tokenization method derived from pan-genome graphs for better segmentation of DNA sequences. Secondly, our pre-training corpus utilizes sparse pan-genomic graphs with a limited number of haplotypes, suggesting that expanding and refining the graph could enhance DeepGene’s future performance. Thirdly, DeepGene currently relies solely on human pan-genomic data, and we propose that incorporating multi-species pan-genomic data could enrich the model with diverse biological vocabularies and language structures. Lastly, while we’ve verified the model’s length extrapolation capability to a certain extent, relying solely on this for longer sequences may be unrealistic, emphasizing the importance of considering necessary modifications such as implementing sparse attention layers to address this limitation effectively.

## 4 Conclusion and Future Work

Existing foundation models for genomics have paid little attention to pan-genomic data, and even when using pangenomic corpora, they are based on multiple sequence alignment. To create an effective foundation model with the understanding of genetic language diversity, we use the Minigraph representation and extract token and positional information from the pan-genomic graph data through a series of operations, including DAG construction, BPE tokenization, and serialization. To endow the model with better length extrapolation capability we select the rotary position embedding as the basis for pre-training.

Finally, we attain competitive performance compared to other baseline models on the GUE and LPD tasks, utilizing less training data, shorter training time, and a significantly smaller model size. DeepGene secures top-2 results in half of the subtasks and reaches the highest average score in GUE, highlighting the potency of language diversity introduced by pan-genome graphs. Moreover, it exhibits superior length extrapolation capability compared to models based on absolute positional encoding, as evidenced by the results in the LPD task.

In future work, we intend to address the issues highlighted in the discussion section: (1) Developing a novel tokenization method based on extensive pan-genome graphs to refine the biological vocabulary used in foundation models; (2) Expanding the pan-genome dataset with a larger number of individuals from the 1000 Genomes Project to deepen our understanding of the human genome; (3) Implementing an attention mechanism with linear time complexity to effectively model extremely long sequences; (4) Conducting pre-training on multi-species genomes to create a more versatile foundation model. With the rapid advancement of genomics, we aim for the techniques proposed to drive the advancement, utilization, and evolution of large-scale language models in genomic research.

## 5 Acknowledgments

We sincerely thank Professor Du Zhenfang, Professor Li Liming, Ms. Sun Jinghan, Mr. Xia Qi, and Ms. Yang Wenjie for their invaluable suggestions on this work.

## Footnotes

1 https://github.com/Zhihan1996/DNABERT_2/

## References

Wardah S. Alharbi and Mamoon Rashid. A review of deep learning applications in human genomics using next-generation sequencing data. Human Genomics, 16(1):26, July 2022. ISSN 1479-7364. doi:10.1186/s40246-022-00396-x. URL 10.1186/s40246-022-00396-x.

Jianxiao Liu, Jiying Li, Hai Wang, et al. Application of deep learning in genomics. Science China Life Sciences, 63 (12):1860–1878, December 2020. ISSN 1674-7305, 1869-1889. doi:10.1007/s11427-020-1804-5. URL https://link.springer.com/10.1007/s11427-020-1804-5.

Zhiqiang Zhang, Yi Zhao, Xiangke Liao, et al. Deep learning in omics: a survey and guideline. Briefings in Functional Genomics, 18(1):41–57, February 2019. ISSN 2041-2649, 2041-2657. doi:10.1093/bfgp/ely030. URL https://academic.oup.com/bfg/article/18/1/41/5107348.

Ashish Vaswani, Noam Shazeer, Niki Parmar, et al. Attention is all you need. Advances in neural information processing systems, 30, 2017. URL https://proceedings.neurips.cc/paper/7181-attention-is-all.

Zhongxiao Li, Elva Gao, Juexiao Zhou, et al. Applications of deep learning in understanding gene regulation. Cell Reports Methods, 3(1), 2023. URL https://www.cell.com/cell-reports-methods/pdf/S2667-2375(22)00289-2.pdf. Publisher: Elsevier.

Gabrielle D. Smith, Wan Hern Ching, Paola Cornejo-Páramo, et al. Decoding enhancer complexity with machine learning and high-throughput discovery. Genome Biology, 24(1):116, May 2023. ISSN 1474-760X. doi:10.1186/s13059-023-02955-4. URL https://genomebiology.biomedcentral.com/articles/10.1186/s13059-023-02955-4.

Hong-Yan Lai, Zhao-Yue Zhang, Zhen-Dong Su, et al. iProEP: A Computational Predictor for Predicting Promoter. Molecular Therapy - Nucleic Acids, 17:337–346, September 2019. ISSN 2162-2531. doi:10.1016/j.omtn.2019.05.028. URL https://www.cell.com/molecular-therapy-family/nucleic-acids/abstract/S2162-2531(19)30161-1. Publisher: Elsevier.

Žiga Avsec, Vikram Agarwal, Daniel Visentin, et al. Effective gene expression prediction from sequence by integrating long-range interactions. Nature Methods, 18(10):1196–1203, October 2021. ISSN 1548-7091, 1548-7105. doi:10.1038/s41592-021-01252-x. URL https://www.nature.com/articles/s41592-021-01252-x.

Yanrong Ji, Zhihan Zhou, Han Liu, et al. DNABERT: pre-trained Bidirectional Encoder Representations from Transformers model for DNA-language in genome. Bioinformatics, 37(15):2112–2120, August 2021. ISSN 1367-4803, 1460-2059. doi:10.1093/bioinformatics/btab083. URL https://academic.oup.com/bioinformatics/article/37/15/2112/6128680.

Hugo Dalla-Torre, Liam Gonzalez, Javier Mendoza-Revilla, et al. The Nucleotide Transformer: Building and Evaluating Robust Foundation Models for Human Genomics, March 2023. URL https://www.biorxiv.org/content/10. 1101/2023.01.11.523679v2. Pages: 2023.01.11.523679 Section: New Results.

Yan Guo, Yulin Dai, Hui Yu, et al. Improvements and impacts of GRCh38 human reference on high throughput sequencing data analysis. Genomics, 109(2):83–90, March 2017. ISSN 0888-7543. doi:10.1016/j.ygeno.2017.01.005. URL https://www.sciencedirect.com/science/article/pii/S0888754317300058.

Rico Sennrich, Barry Haddow, and Alexandra Birch. Neural Machine Translation of Rare Words with Subword Units, June 2016. URL http://arxiv.org/abs/1508.07909. arXiv:1508.07909 [cs].

Eric Nguyen, Michael Poli, Marjan Faizi, et al. HyenaDNA: Long-Range Genomic Sequence Modeling at Single Nucleotide Resolution, November 2023. URL http://arxiv.org/abs/2306.15794. arXiv:2306.15794 [cs, q-bio].

Veniamin Fishman, Yuri Kuratov, Maxim Petrov, et al. GENA-LM: A Family of Open-Source Foundational Models for Long DNA Sequences. bioRxiv, pages 2023–06, 2023. URL https://www.biorxiv.org/content/10.1101/ 2023.06.12.544594.abstract. Publisher: Cold Spring Harbor Laboratory.

Rachel M. Sherman and Steven L. Salzberg. Pan-genomics in the human genome era. Nature Reviews Genetics, 21 (4):243–254, April 2020. ISSN 1471-0064. doi:10.1038/s41576-020-0210-7. URL https://www.nature.com/articles/s41576-020-0210-7. Publisher: Nature Publishing Group.

Jasmijn A. Baaijens, Paola Bonizzoni, Christina Boucher, et al. Computational graph pangenomics: a tutorial on data structures and their applications. Natural Computing, 21(1):81–108, March 2022. ISSN 1567-7818, 1572-9796. doi:10.1007/s11047-022-09882-6. URL https://link.springer.com/10.1007/s11047-022-09882-6.

Liang Zhao, Xiaocheng Feng, Xiachong Feng, et al. Length Extrapolation of Transformers: A Survey from the Perspective of Position Encoding, December 2023. URL http://arxiv.org/abs/2312.17044. arXiv:2312.17044 [cs].

Manzil Zaheer, Guru Guruganesh, Kumar Avinava Dubey, et al. Big bird: Transformers for longer sequences. Advances in neural information processing systems, 33:17283–17297, 2020. URL https://proceedings.neurips.cc/paper/2020/hash/c8512d142a2d849725f31a9a7a361ab9-Abstract.html.

Zhiheng Huang, Davis Liang, Peng Xu, et al. Improve Transformer Models with Better Relative Position Embeddings, September 2020. URL http://arxiv.org/abs/2009.13658. arXiv:2009.13658 [cs].

Francesco Andreace, Pierre Lechat, Yoann Dufresne, et al. Construction and representation of human pangenome graphs. preprint, Bioinformatics, June 2023. URL http://biorxiv.org/lookup/doi/10.1101/2023.06.02.542089.

Phillip E. C. Compeau, Pavel A. Pevzner, and Glenn Tesler. How to apply de Bruijn graphs to genome assembly. Nature Biotechnology, 29(11):987–991, November 2011. ISSN 1546-1696. doi:10.1038/nbt.2023. URL https://www.nature.com/articles/nbt.2023. Publisher: Nature Publishing Group.

Jouni Sirén. Indexing Variation Graphs. In 2017 Proceedings of the Meeting on Algorithm Engineering and Experiments (ALENEX), Proceedings, pages 13–27. Society for Industrial and Applied Mathematics, January 2017. doi:10.1137/1.9781611974768.2. URL https://epubs.siam.org/doi/abs/10.1137/1.9781611974768.2.

Jordan M. Eizenga, Adam M. Novak, Jonas A. Sibbesen, et al. Pangenome Graphs. Annual Review of Genomics and Human Genetics, 21(1):139–162, August 2020. ISSN 1527-8204, 1545-293X. doi:10.1146/annurev-genom-120219-080406. URL https://www.annualreviews.org/doi/10.1146/annurev-genom-120219-080406.

Wen-Wei Liao, Mobin Asri, Jana Ebler, et al. A draft human pangenome reference. Nature, 617(7960):312– 324, May 2023. ISSN 1476-4687. doi:10.1038/s41586-023-05896-x. URL 10.1038/s41586-023-05896-x.

Heng Li, Xiaowen Feng, and Chong Chu. The design and construction of reference pangenome graphs with minigraph. Genome Biology, 21(1):265, December 2020. ISSN 1474-760X. doi:10.1186/s13059-020-02168-z. URL https://genomebiology.biomedcentral.com/articles/10.1186/s13059-020-02168-z.

Joel Armstrong, Glenn Hickey, Mark Diekhans, et al. Progressive Cactus is a multiple-genome aligner for the thousand-genome era. Nature, 587(7833):246–251, 2020. URL https://www.nature.com/articles/ s41586-020-2871-y. Publisher: Nature Publishing Group UK London.

Glenn Hickey, Jean Monlong, Jana Ebler, et al. Pangenome graph construction from genome alignments with Minigraph-Cactus. Nature biotechnology, pages 1–11, 2023. URL https://www.nature.com/articles/ s41587-023-01793-w. Publisher: Nature Publishing Group US New York.

Jianlin Su, Murtadha Ahmed, Yu Lu, et al. Roformer: Enhanced transformer with rotary position embedding. Neurocomputing, 568:127063, 2024. URL https://www.sciencedirect.com/science/article/pii/S0925231223011864. Publisher: Elsevier.

René Dreos, Giovanna Ambrosini, Rouayda Cavin Périer, et al. EPD and EPDnew, high-quality promoter resources in the next-generation sequencing era. Nucleic Acids Research, 41(D1):D157–D164, January 2013. ISSN 0305-1048. doi:10.1093/nar/gks1233. URL 10.1093/nar/gks1233.

